# Detergent-Free Nuclear–Cytoplasmic Fractionation Enables Spatially Resolved PELSA for Enhanced Nuclear Drug Target Identification

**DOI:** 10.64898/2026.04.10.717665

**Authors:** Danni Cai, Kun Zou, Jiayi Wang, Haiyang Zhu, Yanni Ma, Jiayang Yan, Dian Yang, Xiaolei Zhang, Lijuan Zou, Keyun Wang, Mingliang Ye

**Affiliations:** State Key Laboratory of Medical Proteomics, CAS Key Laboratory of Separation Sciences for Analytical Chemistry, National Chromatographic R&A Center, Dalian Institute of Chemical Physics, Chinese Academy of Sciences (CAS), Dalian, 116023, China; Department of Radiation Oncology, The First Affiliated Hospital of Dalian Medical University, Dalian, 116023, China; Department of Radiation Oncology, The Second Affiliated Hospital of Dalian Medical University, Dalian, 116023, China; University of Chinese Academy of Sciences, Beijing, 101408, China

## Abstract

Accurate identification of drug target proteins remain major challenges in proteomics-based target discovery, particularly for low-abundance nuclear proteins that are difficult to detect because of the complexity of whole-cell lysates. Here, we developed a detergent-free nuclear–cytoplasmic fractionation strategy compatible with peptide-centric local stability analysis (PELSA), which markedly improves detection of nuclear drug targets. Using K562 cells, we demonstrated that mild detergent-free fractionation enables high-fidelity nuclear–cytoplasmic separation with minimal cross-contamination. When coupled with PELSA, this workflow significantly increases the number of detected nuclear targets relative to whole-cell analysis. Benchmarking with well-characterized nuclear drugs, including the histone deacetylase inhibitor panobinostat and the RNA polymerase II inhibitor α-amanitin, our results showed improved identification of canonical nuclear targets. Broad profiling of staurosporine target further revealed expanded kinase target coverage by combining the results of nuclear and cytoplasmic fraction, with the CLK family kinases detected exclusively in the nuclear fractions. Additionally, we showed that PELSA can also be performed on intact nucleus level. Collectively, these findings establish detergent-free nuclear–cytoplasmic fractionation–PELSA as a robust and scalable strategy for spatially resolved drug target identification, improving sensitivity for nuclear and low-abundance proteins.

## Introduction

Comprehensive characterization of target engagement are fundamental to drug discovery, mechanism-of-action studies, and translational pharmacology. Proteomics-based approaches have emerged as powerful tools for unbiased target identification, enabling system-level identification of drug–protein interactions without needing prior knowledge of binding partners^1,2^. Among these strategies, proximity- and stability–based proteomic methods have gained increasing attention because they capture drug-induced protein behavior changes in native cellular environments ^3^.

Despite these advances, current proteomics-based target-identification workflows are typically inefficient in detecting drug targets of low abundance. This limitation largely arises from the high complexity and dynamic range of whole-cell lysates, in which abundant cytoplasmic proteins dominate mass spectrometry signals and mask low-abundance targets ^4,5^. Consequently, many biologically important proteins—particularly nuclear proteins such as transcription factors, chromatin regulators, and nuclear enzymes—remain underrepresented or undetected^6,7^. This limitation poses a major challenge for drugs whose mechanisms primarily involve nuclear processes ^8^.

PELSA (PEptide-centric Local Stability Analysis) has recently emerged as a powerful strategy for the identification of drug target proteins at proteome level^9,10^. When a ligand binds to a specific region of a protein, it induces local conformational stabilization at the binding interface. This stabilization effect alters the susceptibility of the affected protein region to proteolytic digestion, forming the basis for binding site determination. Compared to previous limited proteolysis-mass spectrometry (LiP-MS)^11^, PELSA improves target identification sensitivity by 12-fold. However, when applied to whole-cell proteomes, PELSA performance remains constrained by the high proteome complexity that also affect other global target-identification methods^8,12^.

Subcellular fractionation offers an efficient yet underexplored solution to these challenges. By separating nuclear and cytoplasmic components, sample complexity can be substantially reduced, thereby improving detection sensitivity for compartmentalized proteomes^12^. Here, we improved classical nuclear–cytoplasmic fractionation and found that efficient separation can be achieved under mild detergent-free conditions. This approach not only preserves native protein conformation but also enhances the identification of low-abundance targets.

In this study, we integrated detergent-free nuclear–cytoplasmic fractionation with the PELSA workflow to systematically enhance drug target identification. By applying PELSA to isolated nuclear and cytoplasmic proteomes, we observed a substantial increase in reliably identified drug target proteins compared with conventional whole-cell PELSA analysis. Notably, fractionation-assisted PELSA significantly improved the detection of nuclear-localized targets, thereby expanding the detectable target landscape. Overall, this detergent-free nuclear fractionation–assisted PELSA strategy provides a scalable and broadly applicable framework for proteomics-based drug target discovery.

## Results

### Detergent-free nuclear–cytoplasmic fractionation enables efficient compartment separation while preserving protein conformation

Conventional nuclear–cytoplasmic fractionation workflows are typically optimized for maximal protein extraction and therefore often rely on detergent-assisted extraction. However, such conditions are not compatible with PELSA analysis, in which the detergent will denature some proteins and make them loss their ability to binding ligand. To overcome this limitation, we developed a mild, detergent-free nuclear–cytoplasmic fractionation strategy that can preserve native protein conformations while maintaining sufficient protein recovery and proteome-wide identification coverage.

We first evaluated the fractionation efficiency and proteome-level performance of this optimized workflow. Western blot analysis showed strong enrichment of the nuclear marker Lamin B1 in the nuclear fraction and of the cytoplasmic marker α-tubulin in the cytoplasmic fraction, with minimal cross-contamination observed between compartments (Figure 1A), indicating that effective compartment separation can be achieved under detergent-free conditions. Although detergent-free extraction yielded a slightly lower amount of nuclear proteins, this gentler workflow is critical for preserving protein conformation for downstream PELSA analysis.

**Figure 1.**
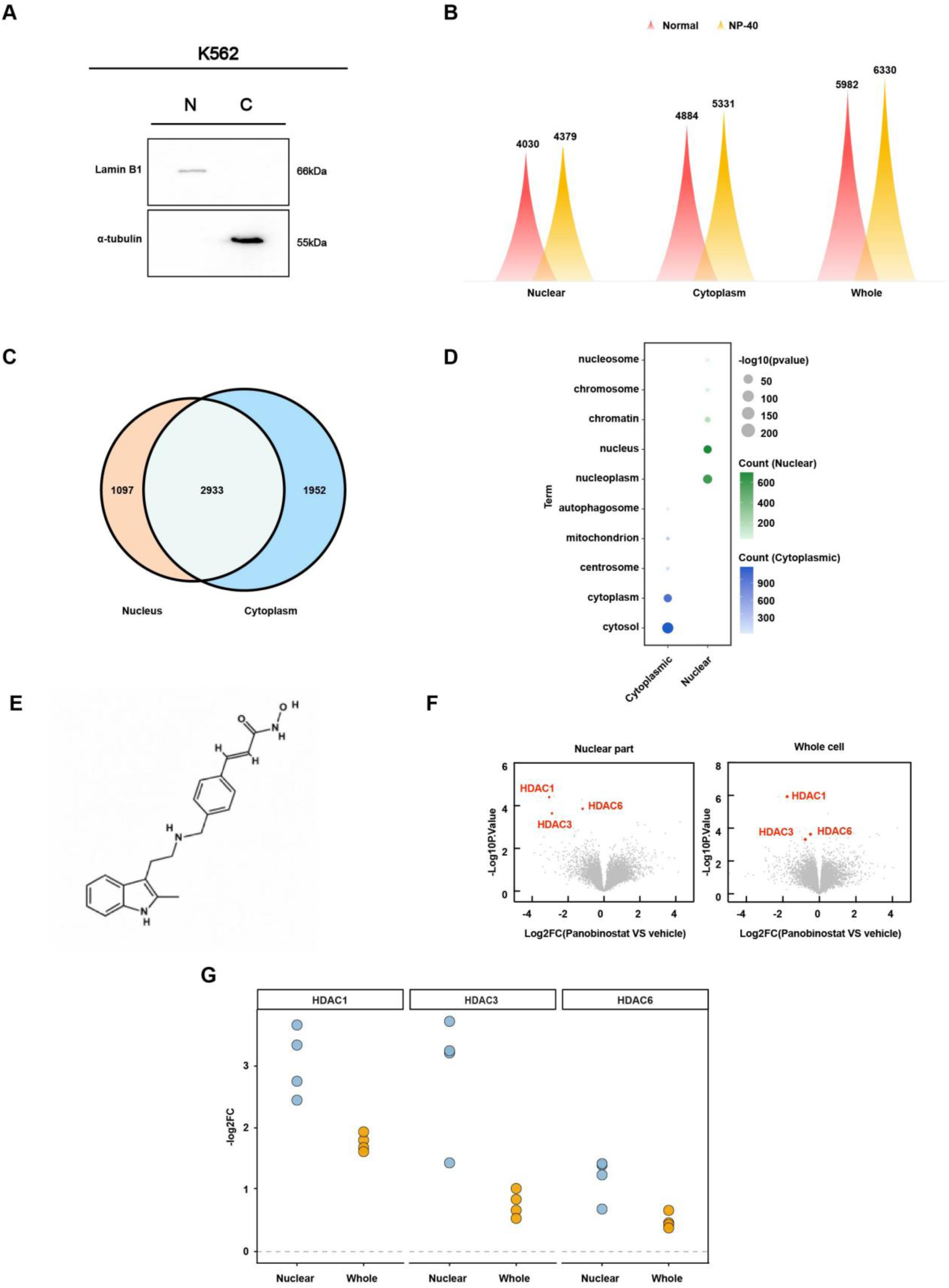
Detergent-free nuclear–cytoplasmic fractionation enables efficient compartment separation and PELSA analysis. (A) Western blot validation of nuclear–cytoplasmic fractionation in K562 cells in the absence of NP-40. Lamin B1 was enriched in the nuclear fraction, whereas α-tubulin was predominantly detected in the cytoplasmic fraction, indicating minimal cross-contamination between compartments. (B) Comparison of the numbers of proteins identified in nuclear, cytoplasmic, or combining results of both two fractions under detergent-free or NP-40-assisted conditions. (C) Venn diagram showing the overlap of proteins identified in nuclear and cytoplasmic fractions under detergent-free conditions. (D) Gene Ontology cellular component enrichment analysis of proteins identified in nuclear and cytoplasmic fractions. Bubble size represents protein counts, and color intensity indicates enrichment significance. (E) Chemical structure of panobinostat. (F) Protein-level volcano plots for comparing ligand-dependent stability changes detected by nuclear-fraction PELSA and whole-cell PELSA following panobinostat treatment. Known HDAC targets are highlighted in red. (G) Quantitative comparison of fold changes (|log_2_FC|) for representative HDAC targets (HDAC1, HDAC3, and HDAC6) identified by nuclear-fraction PELSA and whole-cell PELSA. Each point represents an independent biological replicate.

Proteomic analysis further revealed marked differences in subcellular localization distribution between proteins identified in nuclear and cytoplasmic fractions. The number of proteins identified in each fraction under detergent-free conditions was comparable to that obtained with NP-40-assisted extraction (Figure 1B).1,952 were specific to the cytoplasmic fraction, and 2,933 were shared between the two compartments (Figure 1C), which was comparable to the results obtained using conventional NP-40-assisted extraction (Figure S1). These data demonstrated that efficient nuclear–cytoplasmic separation can be achieved without detergent.

We next examined the extent of compartment-specific proteome enrichment of proteins within subcellular regions. Gene Ontology enrichment analysis consistently showed that nuclear fractions were enriched for nucleus- and chromatin-related terms, whereas cytoplasmic fractions were enriched for cytosolic and metabolism-related proteins (Figure 1D and Figure S2), suggesting strong compartment specificity at the proteome level. Principal component analysis further showed clear separation between nuclear and cytoplasmic samples, with tight clustering of biological replicates (Figure S3), confirming the reproducibility and validity of the fractionation workflow.

Finally, to examine whether the developed detergent-free nuclear–cytoplasmic fractionation method can preserve the native conformations to allow ligand binding. This workflow was coupled to PELSA using the nuclear-targeting HDAC inhibitor panobinostat (Figure 1E). Protein level results showed that compared with the traditional PELSA approach, nuclear-fraction PELSA successfully identified the known HDAC targets of panobinostat (Figure 1F), providing evidence that the detergent-free nuclear fractionation method is fully compatible with PELSA. And slightly larger fold-change was observed for the nuclear-fraction PELSA results, which indicates the nuclear fractionation can improve the sensitivity of nuclear target detection (Figure 1G).

Taken together, these results demonstrate that the detergent-free nuclear–cytoplasmic fractionation workflow enables efficient, reproducible compartment separation while preserving protein native conformation, thereby providing a robust foundation for compartment-resolved PELSA-based drug target identification.

### Detergent-free nuclear-cytoplasmic fractionation–PELSA expands kinase target coverage

To evaluate whether detergent-free nuclear-cytoplasmic fractionation can improve kinase target identification, we compared PELSA results of whole-cell, nuclear, and cytoplasmic samples following treatment with 20 μM staurosporine, a broad-spectrum kinase inhibitor widely used for benchmarking target-identification strategies. Protein-level volcano plots revealed the kinase target distributions across different sample types (Figure 2A). Combining the results from different subcellular fractions, the fractionation-based PELSA workflow quantified much more proteins than conventional whole-cell PELSA (Figure 2B, left), and the number of identified kinase targets, defined using a 75% kinase-specificity threshold, increased by approximately 1.5-fold (Figure 2B, right). These findings indicate that nuclear–cytoplasmic fractionation can expands the detectable subcellular proteome space and thereby increases kinase target identification coverage.

**Figure 2.**
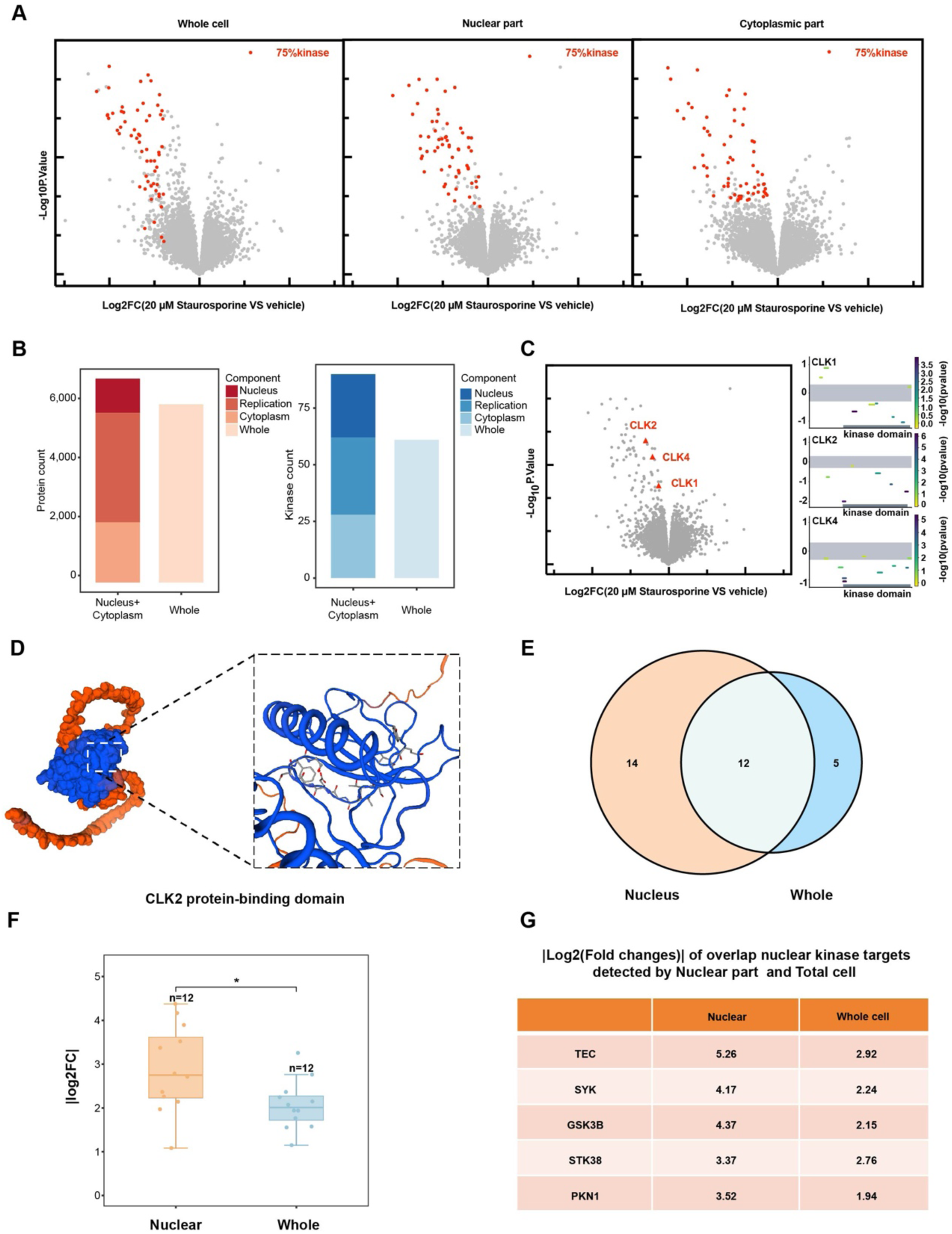
Detergent-free nuclear–cytoplasmic fractionation–PELSA expands kinase target coverage. (A) Protein-level volcano plots showing ligand-dependent stability changes induced by 20 μM staurosporine in whole-cell, nuclear, and cytoplasmic samples. Candidate kinase targets with ≥75% kinase specificity are highlighted in red. (B) Quantitative comparison of proteins and kinase targets identified by fractionation-assisted PELSA and conventional whole-cell PELSA. Left, total quantified proteins in the combined nuclear and cytoplasmic fractions versus whole-cell lysate. Right, candidate kinase targets identified using a ≥75% kinase-specificity threshold. (C) Volcano plot and peptide-level local stability profiles of three CLK family kinases (CLK1, CLK2, and CLK4) specifically detected in the nuclear fraction following staurosporine treatment. (D) Structural mapping of ligand-responsive peptides onto CLK2, highlighting stabilized regions within the protein catalytic domain. The inset shows an enlarged view of the mapped binding region. (E) Venn diagram showing the overlap of kinase targets identified in the nuclear and whole-cell datasets. (F) Comparison of fold changes (|log_2_FC|) for the 12 overlapping kinase targets detected in nuclear and whole-cell samples (n = 12, p < 0.05). (G) Representative overlapping nuclear kinase targets (TEC, SYK, GSK3B, STK38, and PKN1) consistently showed larger fold changes (|log_2_FC|)in nuclear samples than in whole-cell lysates.

Volcano plot analysis showed that three CLK kinases, CLK1, CLK2, and CLK4, exhibited significant ligand-dependent stability changes specifically in the nuclear fraction (Figure 2C, left), whereas these 3 proteins could not be identified in the whole cell lysate and cytoplasmic samples. To examine whether detergent-free fractionation preserves native protein conformation to bind ligands, we analyzed CLK family kinases following staurosporine treatment. Peptide-level local stability mapping demonstrated consistent stabilization patterns across multiple peptides spanning the kinase domains of these CLK proteins (Figure 2C, right). Structural mapping of CLK2 further localized the stability-shifted peptides to the catalytic regions, which is consistent with the binding characteristics of staurosporine, indicating that protein conformation was well preserved during the nuclear fractionation (Figure 2D).

From the whole-cell and nuclear-fraction datasets filtered using the 75% kinase-specificity threshold, we further selected kinase proteins annotated as nuclear-localized in the UniProt database and performed a cross-comparison of the resulting nuclear kinase targets. This analysis identified 12 overlapping kinase targets that were detected in both datasets (Figure 2E). Quantitative analysis showed that these shared targets displayed significantly larger ligand-dependent fold changes (log_2_FC) in the nuclear fraction (Figure 2F). Consistently, representative kinase targets detected in both datasets, including TEC, SYK, GSK3B, STK38, and PKN1, all exhibited larger ligand-dependent fold changes (log_2_FC) in nuclear PELSA than in whole-cell PELSA (Figure 2G).

Collectively, these findings show that detergent-free nuclear–cytoplasmic fractionation–PELSA expands the detectable kinase target space while improving the sensitivity of identifying target proteins.

### Detergent-free nuclear fractionation–PELSA improves the identification of nuclear drug targets

To evaluate whether nuclear fractionation–assisted PELSA enhances detection of nuclear drug targets, we analyzed α-amanitin, a selective RNA polymerase II inhibitor with well-defined nuclear targets (Figure 3A). Protein and peptide-level volcano plots showed that ligand-dependent stability changes were markedly larger in the nuclear fraction than in whole-cell lysates (Figure 3B), indicating that nuclear fractionation substantially improves the sensitivity of nuclear target detection.

**Figure 3.**
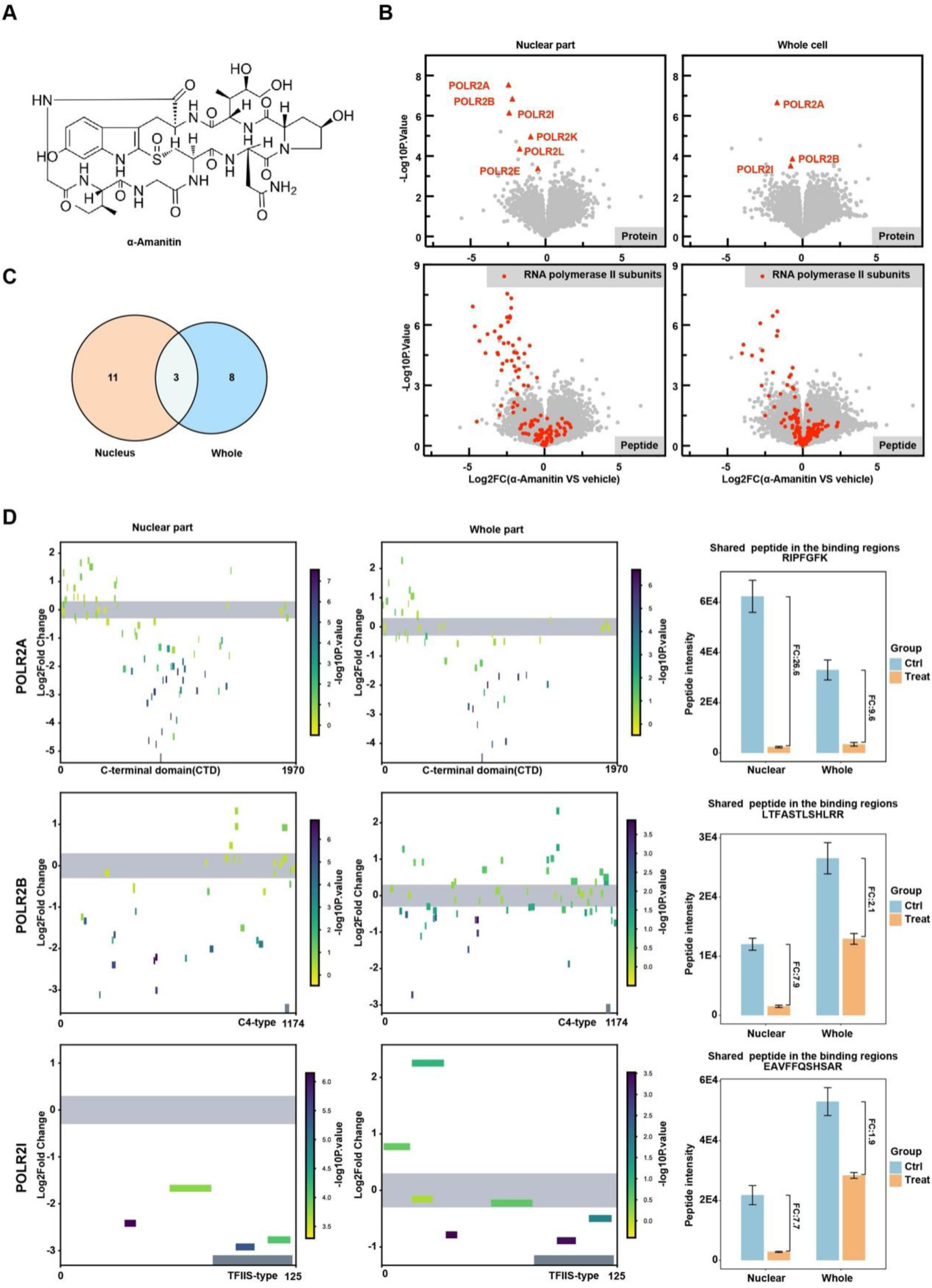
Detergent-free nuclear fractionation–PELSA improves identification of nuclear drug targets. (A) Chemical structure of α-amanitin. (B) Protein- and peptide-level volcano plots comparing ligand-dependent stability changes detected by nuclear-lysate PELSA and whole-cell PELSA following α-amanitin treatment. RNA polymerase II–associated targets are highlighted in red. (C) Venn diagram showing the overlap of significant targets identified in the nuclear and whole-cell datasets using the thresholds −log_10_(p) > 3 and log_2_FC < −0.5. (D) Peptide-level domain-resolved stability analysis of the three shared targets, POLR2A, POLR2B, and POLR2I, for nuclear and whole-cell samples. Bar plots show treatment-induced intensity changes for the shared peptides of each target.

Using stringent filtering criteria (−log_10_(p) > 3 and log_2_FC < −0.5), only three targets overlapped between the nuclear and whole-cell datasets (Figure 3C), whereas a larger number of targets were uniquely identified in the nuclear fraction. These results indicate that nuclear–PELSA substantially expands nuclear target coverage beyond what can be achieved using whole-cell lysates alone.

Quantitative comparison further demonstrated that overlapped peptides exhibited larger fold-change magnitudes in nuclear PELSA than in whole-cell PELSA (Figure 3D). For example, the POLR2A peptide RIPFGFK, located within the target-binding region and detected in both workflows, showed a larger treatment-induced intensity change in nuclear PELSA than in whole-cell PELSA (FC = 26.6 vs 9.6). These findings further suggest that nuclear–PELSA enhances identification sensitivity of nuclear proteins.

### PELSA can also be conducted at the intact-nucleus level

Although the above results showed that nuclear-fraction PELSA markedly improved the sensitivity of nuclear target detection, it is interesting to check whether PELSA can be conducted at the intact-nucleus level, in which the intranuclear protein interaction information is better retained, by considering that intact nuclei might still permit proteolytic access and generation of detectable peptides after the cleavage of nucleoporins. To explore this possibility, we established an alternative workflow in which PELSA was performed directly on intact nucleus without nuclear lysis (Figure 4A). This strategy preserved nuclear integrity while enabling peptide generation from native nuclear proteins.

**Figure 4.**
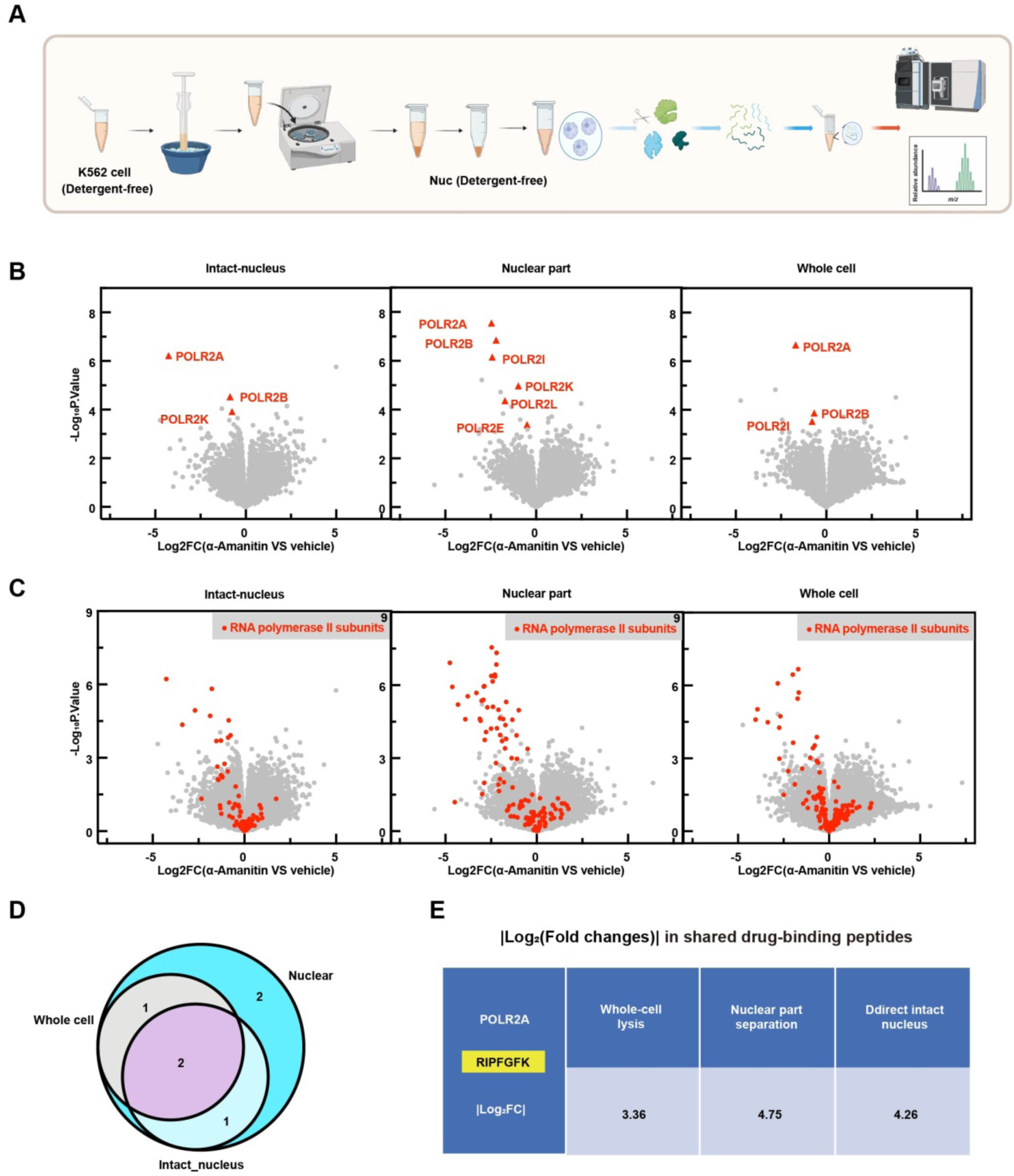
Direct PELSA analysis for intact detergent-free nucleus. (A) Workflow for intact-nucleus PELSA. Following detergent-free nuclear isolation of K562 cells, the intact nuclear fraction was directly subjected to downstream PELSA analysis without nuclear lysis. (B) Protein-level volcano plots for comparing ligand-dependent stability changes detected by intact-nucleus PELSA, nuclear-lysate PELSA, and whole-cell PELSA following α-amanitin treatment. Known RNA polymerase II targets are highlighted in red. (C) Peptide-level volcano plots for comparing intact-nucleus, nuclear-lysate, and whole-cell PELSA following α-amanitin treatment. RNA polymerase II–associated peptides are highlighted in red. (D) Overlap of significant targets identified by intact-nucleus PELSA, nuclear-lysate PELSA, and whole-cell PELSA under the filtering criteria −log_10_(p) > 3 and log_2_FC < −0.5. (E) Quantitative comparison of fold changes (|log_2_FC|) for the shared drug-binding peptide RIPFGFK from POLR2A across the whole-cell lysate, nuclear-lysate, and intact-nucleus workflows.

We then compared this intact-nucleus PELSA workflow with nuclear-lysate PELSA and whole-cell PELSA using α-amanitin. Protein-level volcano plot analysis showed that known nuclear drug targets can also be detected in the direct PELSA analysis of intact nucleus (Figure 4B). At the peptide level, both nuclear-lysate PELSA and intact-nucleus PELSA exhibited more pronounced target-associated signals than whole-cell PELSA (Figure 4C). These findings confirmed the feasibility of using intact nucleus for PELSA analysis.

Using the same filtering criteria (−log_10_(p) > 3 and log_2_FC < −0.5), comparison of the number of target proteins identified by three workflows showed that intact-nucleus PELSA clearly outperformed whole-cell PELSA, while its overall performance was lower than that of nuclear-lysate PELSA (Figure 4D). This suggests that, while direct PELSA on intact nucleus maximally preserves native protein conformation and in situ target-binding states, its overall identification performance is somewhat reduced relative to nuclear-lysate PELSA, which might be caused by the less efficient protein cleavage and lower peptide recovery from certain nuclear proteins or complexes. Quantitative analysis of overlapped drug-binding peptides further showed that both nuclear-lysate PELSA and intact-nucleus PELSA produced larger ligand-dependent stability changes than whole-cell lysate analysis. For example, the POLR2A peptide RIPFGFK exhibited larger stability changes in both nuclear-lysate and intact-nucleus workflows than in whole-cell workflow (Figure 4E).

## Discussion

Proteomics-based drug target identification is often constrained by the complexity of whole-cell lysates and the underrepresentation of low-abundance nuclear proteins^13,14^. In this study, we developed a detergent-free nuclear–cytoplasmic fractionation workflow for peptide-centric local stability analysis (PELSA) to enable spatially resolved target discovery while preserving native protein conformations. This strategy achieved highly reproducible compartment separation with minimal cross-contamination and significantly improved detection of nuclear drug targets compared with the conventional whole-cell PELSA analysis.

The advantage of this workflow was validated through benchmarking with well-characterized nuclear-active compounds. For the HDAC inhibitor panobinostat, nuclear-fraction PELSA produced more pronounced ligand-dependent stability changes than whole-cell PELSA, particularly for canonical HDAC targets such as HDAC1, HDAC3, and HDAC6. For the RNA polymerase II inhibitor α-amanitin, nuclear PELSA identifies approximately twice as many known nuclear target proteins than whole-cell PELSA and thereby demonstrating a substantial expansion of nuclear target coverage. The POLR2A peptide RIPFGFK exhibited larger change in nuclear PELSA than in whole-cell PELSA (FC = 26.6 vs 9.6), suggesting enhanced identification sensitivity for the nuclear proteins. Kinome profiling with staurosporine similarly revealed an expanded detectable target space, with candidate kinase identification increased by approximately 1.5-fold relative to whole-cell analysis. Notably, three CLK family kinases (CLK1, CLK2, and CLK4) were detected exclusively in the nuclear fraction. For 12 kinase targets shared between the nuclear and whole-cell datasets, the nuclear workflow consistently displays overall larger fold changes. Collectively, these findings indicate that reducing proteome complexity through detergent-free subcellular fractionation substantially improves target coverage for nuclear proteins.

Several factors likely contribute to the observed gain in sensitivity. First, nuclear–cytoplasmic fractionation reduces the dynamic range of proteins analyzed within each compartment, allowing low-abundance nuclear proteins to become detectable^15^. Second, detergent-free conditions preserve native protein conformations and ligand-binding properties, which are essential for successful PELSA analysis^16^. Third, compartment-specific enrichment improves the signal-to-noise ratio in mass spectrometric analysis, enabling better detection of ligand-dependent stability changes^17^.

Traditional approaches for drug target identification, including thermal proteome profiling, limited proteolysis–mass spectrometry, and proximity-labeling strategies, have provided important advances but remain constrained by whole-cell proteome complexity ^18^. The detergent-free fractionation-assisted PELSA strategy presented here overcomes this limitation by combining spatial separation with ligand-dependent proteolysis analysis to improve the sensitivity in identifying ligand binding proteins, and this subcellular fractionation workflow should also be beneficial for other drug target identification approaches.

We further showed that PELSA analysis can be conducted directly for intact nucleus and was still able to detect a subset of the ligand-binding proteins observed in nuclear-lysate PELSA. This workflow preserves nuclear structural integrity and the in-situ drug–protein binding state to the greatest extent. However, because protein cleavage is less efficient than in nuclear lysate, its overall performance is slightly lower than that of the nuclear-lysate PELSA workflow. Therefore, nuclear-lysate PELSA enables better target identification, making it more suitable for systematic screening of nuclear drug targets.

Despite these advantages, several limitations should be noted. Nuclear fractionation may not efficiently recover membrane-associated or highly insoluble nuclear complexes^19^. Although detergent-free conditions preserve protein native conformation, extraction efficiency for certain nuclear proteins may still be reduced^20^. In addition, the present study focused primarily on nuclear–cytoplasmic separation, and extension of this strategy to other subcellular compartments, such as mitochondria or chromatin-bound fractions, may require further optimization^21^. Future work could integrate multi-organelle fractionation, single-cell proteomics, or cross-linking strategies to further improve spatial resolution and target characterization^22^.

Overall, this study establishes detergent-free nuclear–cytoplasmic fractionation–PELSA as a robust and scalable strategy for spatial proteomics-based drug target identification. By markedly improving sensitivity for nuclear and low-abundance proteins while preserving native ligand-binding activity, this workflow provides a practical platform for mechanism-of-action studies and drug discovery applications and should be broadly applicable across diverse ligand classes, and stability-based proteomic workflows.

## Method

### Chemicals and Reagents

Dimethyl sulfoxide (DMSO), protease inhibitor cocktail, formic acid (FA), tris(2-carboxyethyl) phosphine (TCEP), 2-chloroacetamide (CAA), ethanol, guanidine hydrochloride, urea, and 4-(2-hydroxyethyl)-1-piperazineethanesulfonic acid (HEPES) were purchased from Sigma-Aldrich (St. Louis, MO, USA). Panobinostat, α-Amanitin, and Staurosporine were obtained from Selleck Chemicals (Houston, TX, USA). Sequencing-grade modified trypsin was purchased from Promega (Madison, WI, USA). HPLC-grade acetonitrile and methanol were obtained from Merck (Darmstadt, Germany).

### Cell Culture

K562 cells (ATCC, CCL-243) were cultured in RPMI-1640 medium (Gibco, USA) supplemented with 10% fetal bovine serum (FBS; Gibco, USA) and 1% penicillin–streptomycin (Beyond Biotechnology, China). Cells were maintained at 37 °C in a humidified incubator with 5% CO₂.

### Detergent-Free Nuclear–Cytoplasmic Fractionation

Subcellular fractionation compatible with PELSA analysis was performed using a detergent-free workflow adapted from classical nuclear extraction strategies^23^ with modifications to preserve native protein conformations and downstream compatibility with stability-based proteomics^24^. Briefly, K562 cells were harvested, followed by addition of cytoplasmic extraction buffer without detergent (Buffer 1: 10 mM HEPES–KOH, pH 7.9; 10 mM KCl; 1.5 mM MgCl₂; protease inhibitor cocktail; H₂O to volume). After the cells were resuspended evenly in Buffer 1 and incubated on ice, they were gently homogenized using a Dounce homogenizer (25 strokes) to disrupt the plasma membrane while preserving nuclear integrity. The lysate was centrifuged at low speed to pellet intact nuclei, and the supernatant was collected as the cytoplasmic fraction. The nuclear pellet was washed twice with Buffer 1 to remove residual cytoplasmic proteins and then resuspended in nuclear extraction buffer without detergent (Buffer 2: 20 mM HEPES–KOH, pH 7.9; 2 mM MgCl₂; 420 mM NaCl; 1 mM DTT; protease inhibitor cocktail). Samples were incubated with gentle rotation at 4 °C for 1 h to release nuclear proteins, followed by centrifugation at 20,000 × g for 10 min to obtain the nuclear protein fraction.

The resulting cytoplasmic and nuclear extracts were subjected to Western blotting and LC–MS/MS analysis to evaluate fractionation efficiency and compatibility with downstream PELSA workflows.

### Peptide-centric local stability assay analysis

PELSA experiments were performed essentially according to the recently described peptide-centric local stability assay workflow, with minor modifications^9^. K562 cells were cultured under standard conditions and harvested at approximately 80% confluence. Cells were washed three times with ice-cold PBS, collected by centrifugation, and stored at −80 °C until use. Cell pellets were resuspended in PBS containing 1% protease inhibitor cocktail and lysed by three freeze–thaw cycles using liquid nitrogen and a 37 °C water bath. Lysates were clarified by centrifugation at 1000× g for 10 min at 4 °C, and protein concentrations were determined using a Pierce 660 nm protein assay and adjusted to 1 mg mL⁻¹. Aliquots of lysate (50 µL each) were incubated with compound or equal-volume DMSO control for 30 min at 25℃ (n = 4 per group). Samples were then subjected to limited proteolysis with trypsin at an enzyme-to-protein ratio of 1:2 (w/w) at 37 °C for 1 min, and digestion was terminated with HEPES buffer containing 8 M guanidine hydrochloride. Proteins were reduced with 10 mM TCEP and alkylated with 40 mM CAA at 95 °C for 5 min. Peptides were collected using 10 kDa ultrafiltration, washed with HEPES buffer (pH 8.2), acidified with 1% trifluoroacetic acid, desalted using HLB tips, lyophilized, and stored at −20 °C prior to LC–MS/MS analysis with data-independent acquisition (DIA).

### LC-MS/MS analysis

Raw DIA files from PELSA experiments were processed in Spectronaut (version 18.1, Biognosis) using the directDIA workflow without generation of a spectral library. The use of Spectronaut directDIA and related DIA informatics workflows has been benchmarked in recent comparative studies^25^. Spectra were searched against the UniProt reference proteome (20,350 entries, downloaded 202006). Trypsin was specified as the digestion enzyme with up to two missed cleavages allowed. Carbamidomethylation of cysteine was set as a fixed modification, and methionine oxidation and protein N-terminal acetylation were included as variable modifications. Peptide quantification was normalized using the local normalization algorithm, and all other parameters were maintained at default settings. Peptide intensity values (“PEP.Quantity”) and positional information (“PEP.PeptidePosition”) exported from Spectronaut were used for downstream analyses.

### Statistical Analysis

Statistical Analysis Raw output files from Spectronaut were analyzed using PELSA-Decipher 1.6.0.0 (software available at https://github.com/DICP-1809/PELSA-Decipher)^26^. In the DA-generated report, log_2_FC values represent the magnitude of stability changes, whereas −log_10_(p value) indicates statistical significance. ProLSA was used to export local stability maps of ligand-binding regions for further analysis.

### Western blotting

Protein concentrations of nuclear and cytoplasmic fractions were determined by BCA assay. Equal amounts of protein (20μg) were resolved by SDS–PAGE and transferred onto PVDF membranes. Membranes were blocked in TBST containing 5% non-fat milk for 1 h, followed by incubation with primary antibodies overnight at 4 °C. After washing, membranes were incubated with HRP-conjugated secondary antibodies and detected by enhanced chemiluminescence. Lamin B1 and β-tubulin were used as loading controls for nuclear and cytoplasmic fractions, respectively.

### Statistics and reproducibility

All PELSA experiments were performed with 4 biological replicates. No statistical methods were used to predetermine sample size. Samples and cells were randomly assigned to experimental groups. Investigators were not blinded to sample identity, as data were generated using objective quantitative measurements that minimized subjective bias. Unless otherwise specified, all collected data were included in the analysis.

### Bioinformatic analysis

Bioinformatic analyses were performed in R (v4.3.1). Principal component analysis (PCA) was conducted using the prcomp function to evaluate overall variation and clustering among samples. Gene Ontology (GO) enrichment analysis was performed using the cluster Profiler package (enrich GO function) with annotation from the org.Hs.eg.db database. Data visualization was performed using ggplot2.

## ASSOCIATED CONTENT

### Supporting Figures

Supplementary Figures including overlap of proteins identified in NP-40-assisted nuclear and cytoplasmic fractions (Figure S1), KEGG pathway enrichment analysis of detergent-free nuclear and cytoplasmic proteomes(Figure S2), Principal component analysis (PCA) showing clear separation between detergent-free nuclear and cytoplasmic samples(Figure S3).

## AUTHOR INFORMATION

### Corresponding Author

Mingliang Ye − State Key Laboratory of Medical Proteomics, CAS Key Laboratory of Separation Sciences for Analytical Chemistry, National Chromatographic R&A Center, Dalian Institute of Chemical Physics, Chinese Academy of Sciences(CAS), Dalian 116023, China; University of Chinese Academy of Sciences, Beijing 100049, China;

Keyun Wang− State Key Laboratory of Medical Proteomics, CAS Key Laboratory of Separation Sciences for Analytical Chemistry, National Chromatographic R&A Center, Dalian Institute of Chemical Physics, Chinese Academy of Sciences(CAS), Dalian 116023, China; University of Chinese Academy of Sciences, Beijing 100049, China;

Lijuan Zou - Department of Radiotherapy Oncology, The Second Affiliated Hospital of Dalian Medical University, Dalian, 116023, China;

## Author Contributions

K.W. and M.Y. conceived the study and revised the manuscript. D.C. performed the experiments, analyzed the data, and wrote the manuscript. K.Z. participate in some experiments and data analysis and collation. J.W., H.Z. and Y.M. assisted with methodology optimization and data analysis. J.Y., D.Y. and X.Z. contributed to discussion and validation of the experimental results. L.Z. participated in discussion of the results and manuscript revision. K.W. and M.Y. acquired funding. All authors approved the final version of the manuscript.

## Notes

The authors declare no competing financial interest.

## Acknowledgments

We acknowledge the financial supports from the National Key Research and Development Program of China (grant no. 2021YFA1302600 to M.Y., 2024YFA1306000 to K.W.), the National Natural Science Foundation of China (grant nos. 22437007 to M.Y., 22137002 to K.W.), Dalian Science and Technology Innovation Fund (grant no. 2023JJ11CG006 to M.Y.), the Innovation Program of Science and Research from the DICP, CAS (grant no. DICP I202531 to M.Y., DICP I202510 to K.W.), United Foundation for Dalian Institute of Chemical Physics Chinese Academy of Sciences and the Second Hospital of Dalian Medical University (DMU-2&DICP UN202303 to L.Z.).Dalian Science and Technology Talent Innovation Support Plan (2024RY006 to K.Z.).

---

**Scheme 1.**
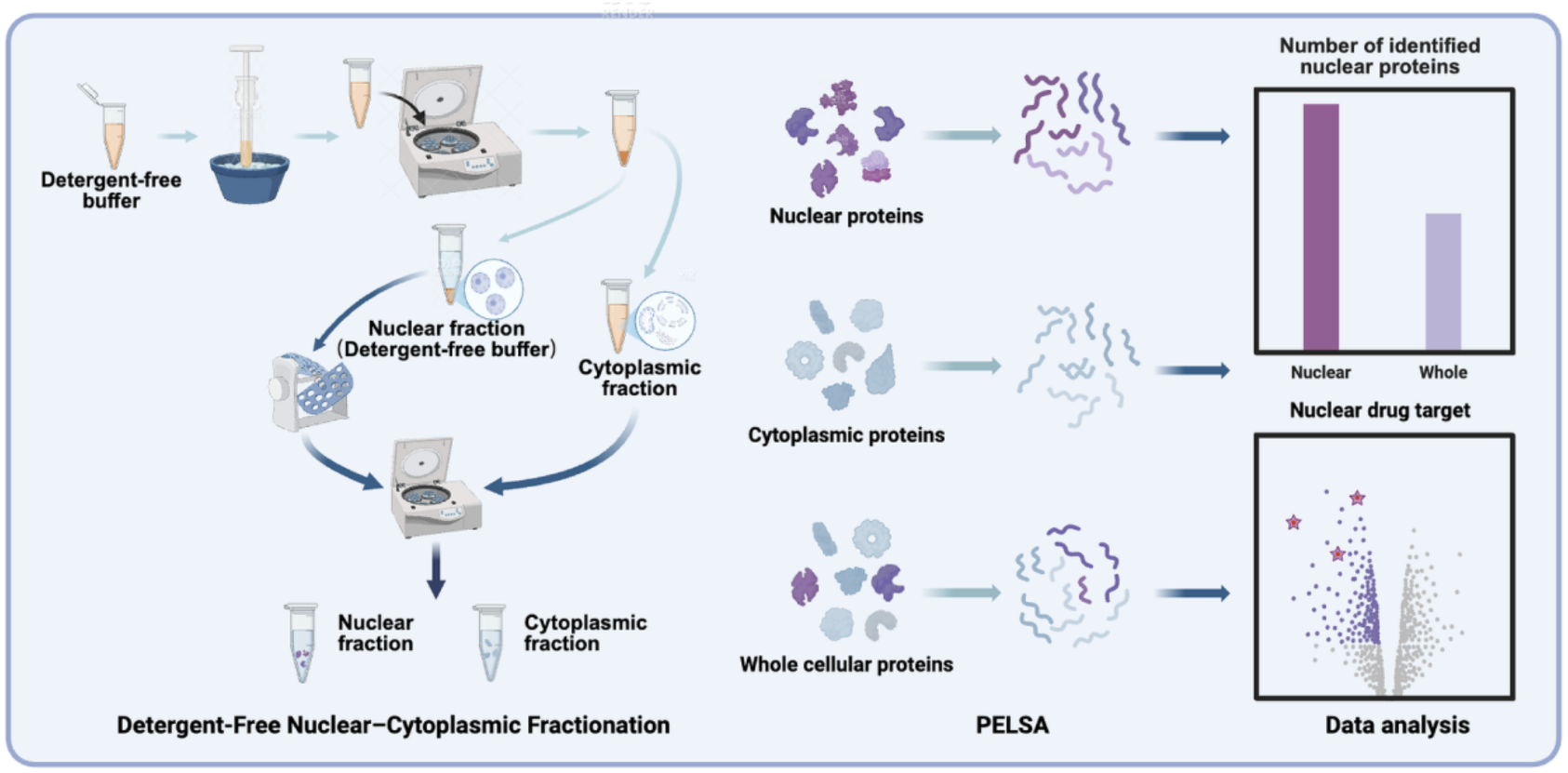
Detergent-free nuclear–cytoplasmic fractionation-PELSA workflow. Cells were gently lysed with buffer without detergent to isolate intact nuclear and cytoplasmic fractions while preserving protein activity, followed by PELSA to identify drug targets.

