## Supporting Information for "Detergent-Free Nuclear–Cytoplasmic Fractionation Enables Spatially Resolved PELSA for Enhanced Nuclear Drug Target Identification"

**Supplementary Figures**

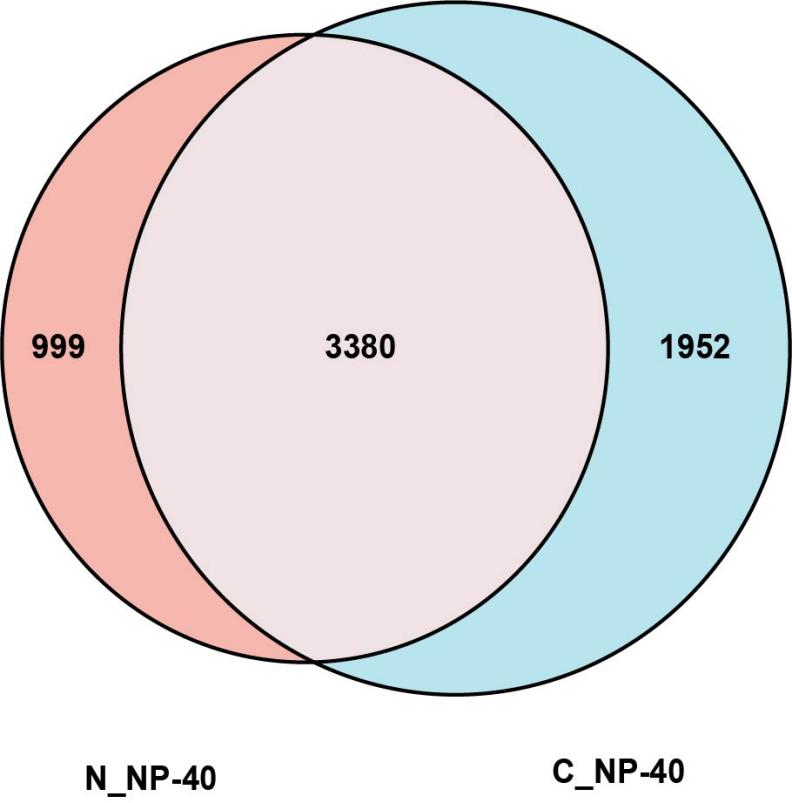

**Figure S1.** Overlap of proteins identified in NP-40-assisted nuclear and cytoplasmic fractions.

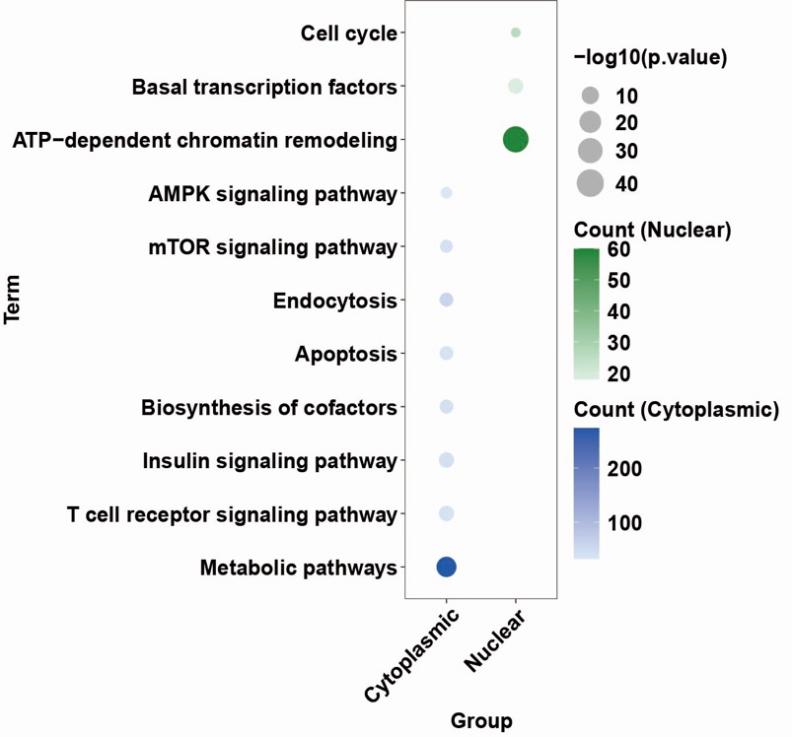

**Figure S2.** KEGG pathway enrichment analysis of detergent-free nuclear and cytoplasmic proteomes.

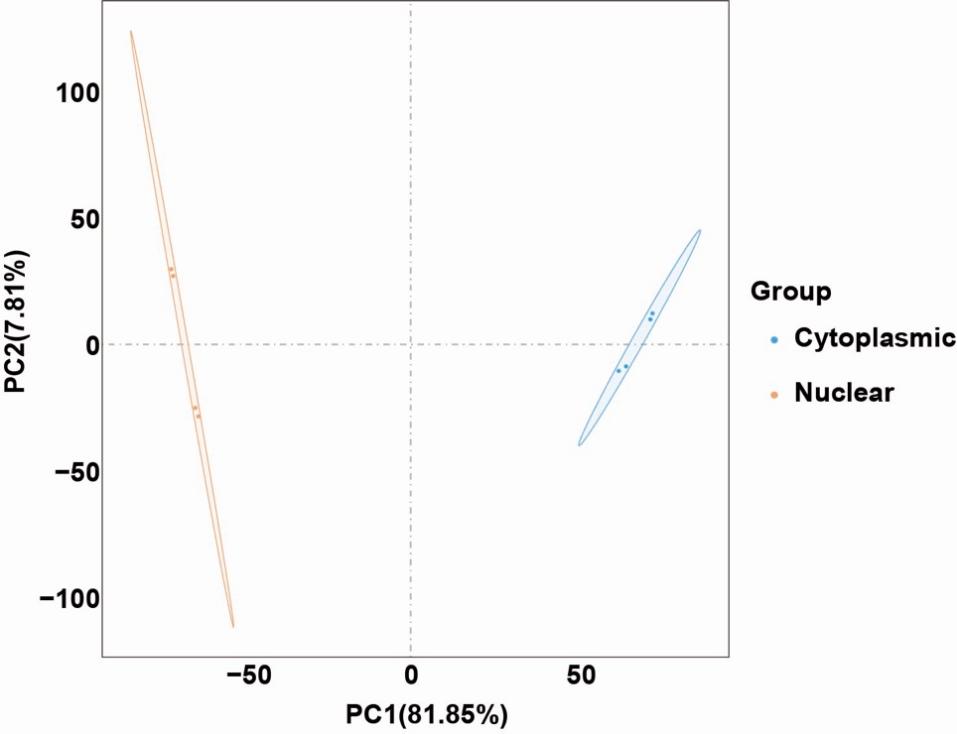

**Figure S3.** Principal component analysis (PCA) showing clear separation between detergent-free nuclear and cytoplasmic samples.
